# The Combinatorial Capacity and Robustness of Hierarchical Concept Coding in the Human Medial Temporal Lobe

**DOI:** 10.64898/2026.02.27.708650

**Authors:** Lihong Cao

## Abstract

The human brain encodes a virtually infinite repertoire of semantic concepts using a finite number of neurons, a feat that defies the capacity limits of classical attractor networks. While “Concept Cells” in the medial temporal lobe (MTL) exhibit extreme sparsity, the information-theoretic principles governing their collective robustness remain elusive. Here, we present a hierarchical coding framework that resolves this paradox. We rigorously prove that while uniform sparse coding is bound by a polynomial “Exclusion Volume” limit—leading to catastrophic interference—a hierarchically partitioned architecture unlocks an exponential combinatorial capacity. We demonstrate that this “locally dense, globally sparse” topology mirrors the distance-dependent connectivity of the hippocampal CA3 region. Crucially, we derive a “Supply-Demand” theoretical model for Cognitive Reserve, which quantitatively predicts the “Silent Phase” of neurodegeneration and the mathematical inevitability of the clinical “Cliff Edge” collapse in Alzheimer’s Disease. Furthermore, our findings identify the lack of topological sparsity as the root cause of catastrophic forgetting in artificial neural networks, offering a blueprint for next-generation, biologically plausible AI architectures.

## 1 Introduction

The human brain possesses an extraordinary capacity to encode a virtually infinite repertoire of semantic concepts—from “Jennifer Aniston” to “Quantum Mechanics”—using a finite number of neurons. The discovery of “Concept Cells” in the medial temporal lobe (MTL) [1], which fire selectively to specific abstract entities, suggests that the brain employs a highly sparse coding scheme [2, 3]. However, the information-theoretic principles enabling this massive storage remain poorly understood.

A fundamental paradox exists between **biological observation** and **computational theory**. Classical attractor models, such as Hopfield networks [4], suffer from severe capacity limitations due to “crosstalk” interference, scaling linearly with network size (*C* ≈ 0.14*N*) [5]. Even standard extensions to sparse coding [6] face a “Curse of Dimensionality”: as the number of stored patterns grows, the **“Exclusion Volume”**—the state space required to prevent collision—expands geometrically, inevitably leading to a “Jamming Limit” where memory retrieval becomes unreliable.

Here, we propose that the brain overcomes this barrier through a **Hierarchical Concept Coding** architecture. By organizing semantic representations into “locally dense, globally sparse” manifolds, the system undergoes a topological phase transition. We rigorously prove that this hierarchy unlocks a **Combinatorial Capacity** that scales exponentially, effectively circumventing the polynomial bounds of uniform coding.

Beyond information theory, this framework offers a unified mechanistic explanation for critical biological and clinical phenomena:

- **Anatomical Fidelity:** It predicts the clustered, distance-dependent connectivity observed in the hippocampal CA3 region [7] and the formation of memory engrams [8].
- **Cognitive Reserve in Dementia:** We derive a “Supply-Demand” model that explains the “Silent Phase” of Alzheimer’s Disease—where patients maintain function despite significant neural loss—and predicts the mathematical inevitability of the sudden “Cliff Edge” collapse [9, 10].
- **Blueprint for Robust AI:** Finally, our results suggest that the catastrophic forgetting and adversarial brittleness plaguing current Artificial Intelligence (AI) [11] stem from the lack of topological partitioning, offering a theoretical blueprint for nextgeneration brain-inspired architectures.

## 2 Theoretical Framework

### 2.1 Parameterization of the Neural Space

We model the Medial Temporal Lobe (MTL) activity using a binary state space. Let the neural representation space be a binary hypercube, consistent with principles of Sparse Distributed Memory [12] ℋ= {0, 1} ^*N*^, where *N* represents the total pool of neurons available for concept encoding. Based on anatomical estimates of the human MTL, we set *N* ≈ 10^7^.

A single semantic concept 𝒞 is represented by a Cell Assembly, defined as a binary vector of constant Hamming weight *M* (number of active neurons), where *M* ≪ *N*. Empirical recordings of “Concept Cells” suggest *M* ≈ 10^3^ ~ 10^4^, corresponding to a sparsity level of 0.01% ~ 0.1%.

### 2.2 The Overlap Constraint Problem

The fundamental limit of memory capacity is determined by the maximum allowable interference between concepts. We define the overlap *λ*(*c*_*i*_, *c*_*j*_) = |*c*_*i*_ ∩ *c*_*j*_| as the number of shared active neurons between two distinct concepts *c*_*i*_ and *c*_*j*_. To ensure reliable retrieval, the system must satisfy specific overlap constraints. We contrast two distinct topological regimes:

#### 2.2.1 Regime I: Uniform Random Coding (𝒰)

In this “flat” architecture, the neural pool is treated as a single homogeneous graph. Any pair of stored concepts must satisfy a strict global overlap threshold *K*:

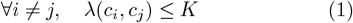

This model assumes that the brain enforces a universal orthogonality constraint across all stored memories. While mathematically simple, it imposes a severe restriction: every new memory consumes “exclusion volume” in the global high-dimensional space.

#### 2.2.2 Regime II: Hierarchical Coding (ℋ)

Reflecting the anatomical structure of the cortex, we partition the total population *N* into *G* functional subspaces (or “macro-columns”), each of size *N*_*loc*_. The coding follows a Dual-Constraint Protocol:

- **Global Separation (***K*_*inter*_): Concepts belonging to different subspaces must be strictly orthogonal or minimally overlapping.

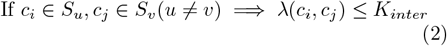

This acts as a “firewall” between distinct semantic categories.
- **Local Tolerance (***K*_*intra*_): Concepts within the same subspace are permitted a relaxed overlap threshold.

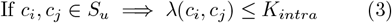

Crucially, we set *K*_*intra*_ ≫ *K*_*inter*_. This relaxation allows for high-density packing of related concepts (e.g., “Jennifer Aniston” and “Lisa Kudrow”) within a local manifold, without compromising the global separability of unrelated concepts.

## 3 Results

### 3.1 Theorem 1: The Limits of Uniform Coding

To quantify the advantage of hierarchy, we first establish the baseline capacity of a non-hierarchical (“flat”) system. In this regime, the entire neural population *N* serves as a single coding pool, and any pair of concepts must satisfy a strict global overlap constraint *λ*(*c*_*i*_, *c*_*j*_) ≤ *K*.

#### Theorem 1

(Uniform Capacity Bounds). *For a uniform coding scheme* 𝒰 (*N, M, K*), *the capacity C*_*u*_ *is bounded by:*

***Upper Bound (Johnson Bound)***

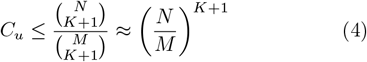

***Lower Bound (Gilbert-Varshamov)***

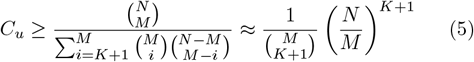

*Proof provided in Methods*.

This theorem reveals a critical scaling limitation: the capacity *C*_*u*_ grows polynomially with *N*, with the exponent determined strictly by the global constraint *K*. For a biologically realistic sparse code where interference must be minimized (e.g., *K* = 2), the capacity is severely throttled. Specifically, the number of forbidden states around each concept scales as 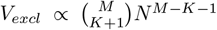. While the total state space grows as *N* ^*M*^, the relative capacity *C* ≈ Ω*/V*_*excl*_ is constrained to scale as *N* ^*K*+1^.This reveals the ‘Curse of Dimensionality’ in uniform coding: to maintain strict separation (low *K*), the system is mathematically forced to waste the vast majority of the high-dimensional combinatorial space, resulting in a storage efficiency that is polynomial rather than exponential.

### 3.2 Theorem 2: Hierarchical Capacity Expansion

In contrast, the hierarchical model partitions the space into subspaces of size *N*_*loc*_, allowing for a relaxed intraclass constraint *K*_*intra*_ ≫ *K*_*inter*_.

#### Theorem 2

(Hierarchical Capacity). *For a hierarchical system* ℋ (*N, N*_*loc*_, *M*, 𝒦), *the capacity C*_*h*_ *scales as:*

***Asymptotic Bound***

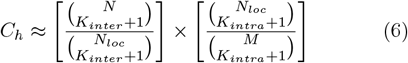

*Which simplifies logarithmically to:*

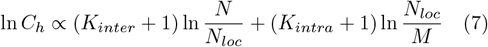

### 3.3 The Gain Ratio

Comparing Theorem 1 and Theorem 2, we derive the fundamental **Gain Ratio** Γ = *C*_*h*_*/C*_*u*_. Assuming the uniform baseline is constrained by the strict inter-class requirement (*K* = *K*_*inter*_):

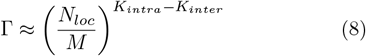

Since *N*_*loc*_ ≫ *M* and *K*_*intra*_ *> K*_*inter*_, this ratio represents an exponential amplification. For instance, with *N*_*loc*_ = 10^5^, *M* = 10^3^, relaxing the local constraint from 2 to 20 yields a gain factor of ~ 100^18^. This proves that the hierarchical architecture is not merely an anatomical detail but a mathematical necessity for high-capacity storage.

### 3.4 The Topological Phase Transition

Our numerical analysis (Figure 2) reveals a fundamental phase transition in neural coding strategies, governed by the trade-off between combinatorial freedom and packing density.

**Figure 1.**
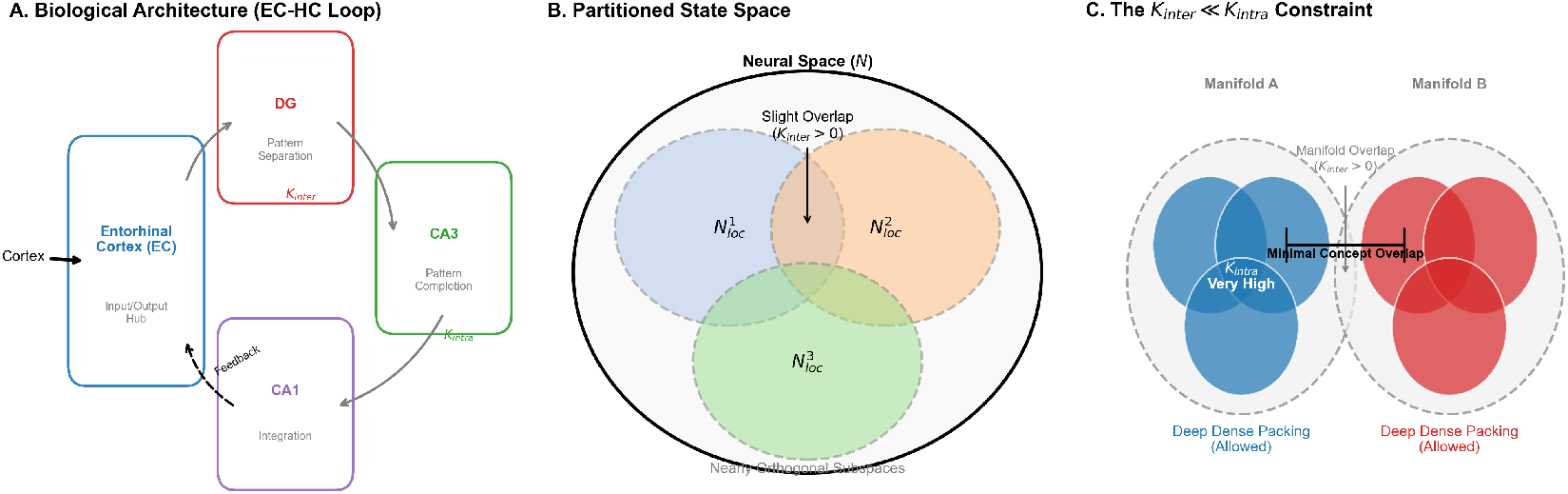
Schematic of the Hierarchical Concept Coding Model. **(A) The EC-Hippocampal Loop:** Sensory inputs enter the Entorhinal Cortex (EC). The circuit proceeds to the Dentate Gyrus (DG), which performs *Pattern Separation* (enforcing *K*_*inter*_), then to CA3, which enables *Pattern Completion* via recurrent connections (supporting *K*_*intra*_), and finally to CA1. A crucial feedback loop (dashed line) returns integrated information to the EC/Neocortex. (B)**Partitioned State Space:** The neural space *N* is modeled as a collection of nearly orthogonal functional subspaces (*N*_*loc*_). The slight overlap indicates that distinct semantic manifolds are weakly coupled (*K*_*inter*_ *>* 0).(C)**The Coupling Asymmetry (***K*_*inter*_ ≪ *K*_*intra*_): *Inside Manifolds:* Concepts (colored spheres) exhibit **Deep Overlap** (*K*_*intra*_ High), mirroring CA3’s auto-associative density. *Between Manifolds:* Distinct concepts maintain **Minimal Overlap** (*K*_*inter*_ Low), mirroring DG’s orthogonalization function.

**Figure 2.**
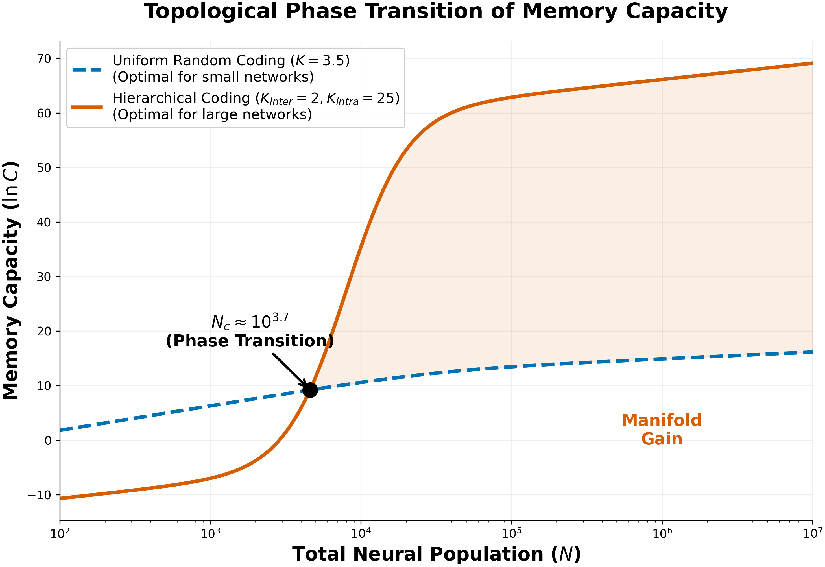
Topological phase transition of memory capacity. The plot illustrates the scaling of logarithmic capacity (ln *C*) as a function of neural population size (*N*). **(Blue Dashed Line)** Uniform Random Coding (*K* = 3.5). In small networks, this strategy is optimal as it utilizes the full combinatorial entropy of the unpartitioned space. However, as *N* exceeds 10^4^, the curve saturates due to the **“Hamming Jamming Limit”**—the exponential rise in collision probability in high-dimensional space. **(Red Solid Line)** Hierarchical Coding (*K*_*inter*_ = 2, *K*_*intra*_ = 25). At small scales (*N <* 10^4^), capacity is suppressed by the **“Structural Overhead”** (wiring cost) of maintaining hierarchy. However, beyond the critical point *N*_*c*_ ≈ 10^4^, the hierarchical architecture effectively “resets” local density, bypassing the jamming limit and maintaining a linear power-law scaling. The shaded region represents the **Manifold Gain**, quantifying the biological advantage of the mammalian cortex over simple invertebrate ganglia.

#### The Small-Scale Regime (*N <* 10^4^)

For neural populations below the critical threshold *N*_*c*_ ≈ 10^4^, the Uniform Coding strategy (Blue Line) proves superior. At this scale, the “curse of dimensionality” has not yet set in. A flat, fully connected network maximizes the combinatorial entropy 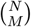 without incurring the overhead of structural partitioning. Conversely, imposing a hierarchical structure on a small network (e.g., *N* = 10^3^) introduces a “wiring tax”—the cost of enforcing empty space (*K*_*inter*_) between nonexistent manifolds—resulting in suboptimal capacity. This explains why simple invertebrate ganglia typically exhibit distributed, non-hierarchical connectivity.

#### The Jamming Limit and Transition

As the system scales to mammalian proportions (*N >* 10^5^), the Uniform model hits a hard geometric barrier: the **“Hamming Jamming Limit.”** The exclusion volume of each concept grows as *N* ^*K*+1^, rapidly filling the available state space. The blue curve saturates (Fig. 2), indicating that adding more neurons yields diminishing returns due to the exponential increase in collision probability.

#### The Large-Scale Regime (*N >* 10^4^)

The Hierarchical model (Red Line) avoids this saturation. By decomposing the global space into local manifolds *N*_*loc*_, the system effectively “resets” the collision counter within each subspace. Although it pays an initial structural cost, it gains access to the relaxed intra-class constraint (*K*_*intra*_ ≈ 25). Beyond *N*_*c*_, this local gain overwhelms the global penalty, allowing capacity to scale linearly with *N* (power-law scaling). The divergence between the two curves at *N* = 10^7^ (human MTL scale) spans orders of magnitude, mathematically validating that hierarchy is an obligatory adaptation for large-scale semantic memory.

### 3.5 Geometric Interpretation of Packing Efficiency

To understand the physical mechanism driving the phase transition, we visualized the high-dimensional Hamming space packing in two dimensions (Figure 3).

**Figure 3.**
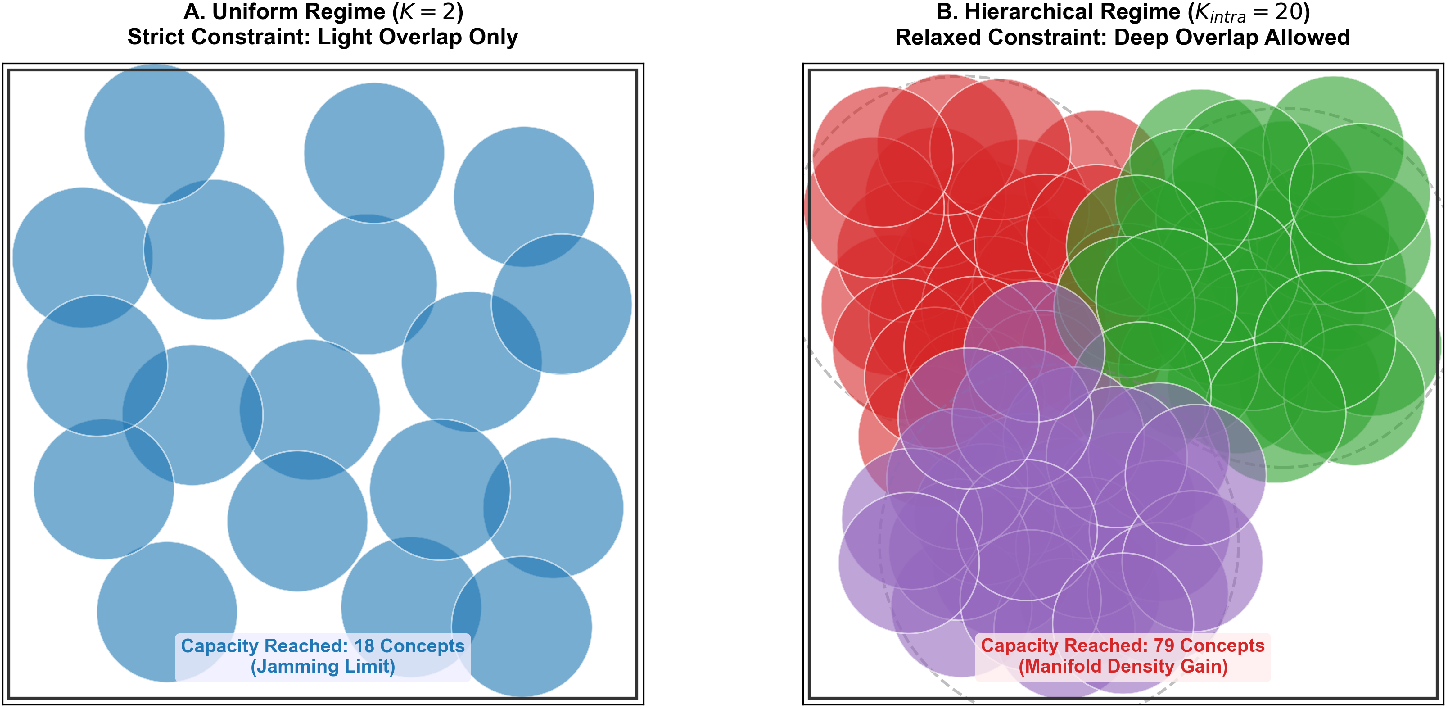
Visualizing the geometry of capacity limits in Hamming space. **(A) Uniform Regime:** Concepts (blue circles) act as quasi-hard spheres distributed across the neural space. The strict global constraint (*K* = 2) creates a rigid exclusion zone around each concept. The system reaches a “Jamming Limit” where the space is saturated, preventing the storage of new memories despite the existence of empty interstitial voids. **(B) Hierarchical Regime:** The neural space is organized into functional manifolds (dashed boundaries). Within each manifold, the relaxed constraint (*K*_*intra*_ = 20) allows for “Deep Overlap” (dense clustering of red/green/purple circles). This architecture avoids global jamming by confining high-density interference locally, effectively unlocking the hyper-volume of the state space.

In the **Uniform Regime** (Fig. 3A), the strict global constraint (*K* ≈ *M/*10) forces concepts to behave like “hard spheres” with large exclusion radii. While slight overlap is mathematically permitted, the system rapidly encounters a **“Jamming Limit.”** At this point, the free volume is fragmented into interstitial voids that are too small to accommodate new concepts without violating the exclusion boundaries of existing ones. This explains the saturation observed in the blue curve of Figure 2.

In contrast, the **Hierarchical Regime** (Fig. 3B) circumvents this jamming by partitioning the space into semantic manifolds. Within each manifold, the relaxed constraint (*K*_*intra*_ ≈*M/*2) allows concepts to behave as “soft spheres,” permitting deep clustering. This **“Deep Stacking”** capability allows the system to utilize the exponential volume of the sphere’s interior, rather than just its surface, resulting in the orders-of-magnitude capacity gain derived in Theorem 2.

### 3.6 Capacity Surplus and Threshold Failure

We define the *Cognitive Reserve* as the margin between the brain’s theoretical storage supply (*C*_*supply*_) and the biological demand (*C*_*req*_).

- **Uniform Failure (Blue Line):** As shown in Theorem 1, uniform coding hits the “Jamming Limit” early. In Fig. 4A, this creates a **Structural Deficit** (*C*_*supply*_ *< C*_*req*_), explaining why non-hierarchical networks cannot support high-dimensional human cognition.
- **Hierarchical Surplus (Red Line):** The hierarchical architecture generates a massive capacity surplus (*C*_*supply*_ ≈ 50 × *C*_*req*_). This surplus acts as a buffer. Despite the exponential decay of theoretical capac-ity due to neurodegeneration (*α* ≈ 18), the observed function remains stable at *C*_*req*_ (Green Zone) until the surplus is exhausted.
- **The Cliff Edge:** The transition from asymptomatic pathology to dementia occurs precisely when the supply curve intersects the demand threshold (*f*_*crit*_ ≈ 20%), triggering a catastrophic functional collapse.

**Figure 4.**
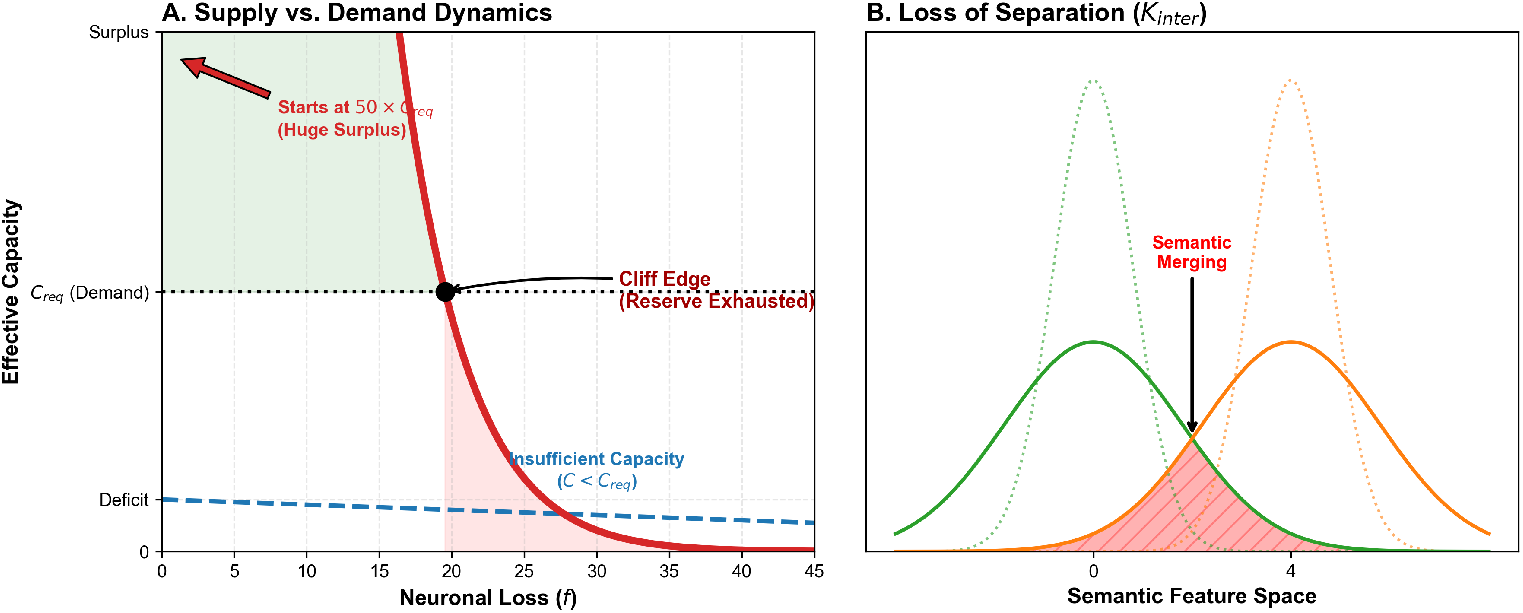
Cognitive Reserve as a Supply-Demand Problem. **(A) The Threshold of Collapse**. The plot is scaled such that the biological cognitive demand (*C*_*req*_) is centered (dotted line). *Uniform Model (Blue):* Suffers from a structural deficit (*C < C*_*req*_), rendering it inviable for complex cognition. *Hierarchical Model (Red):* Starts with a massive theoretical surplus (extending far above the chart). This surplus provides a “Safety Zone” (Green) where neurodegeneration is clinically silent. The “Cliff Edge” (Black Arrow) marks the precise moment when neuronal loss reduces the theoretical supply below the biological demand, resulting in sudden functional failure. **(B) Mechanism of Semantic Merging**. Schematic representation of concept probability density in semantic space. In a healthy brain (dotted lines), strict *K*_*inter*_ constraints maintain sharp boundaries between categories (e.g., Animals vs. Tools). In AD pathology (solid lines), the loss of inhibitory interneurons broadens the tuning curves. The resulting overlap area (Red Shaded Region) represents the mathematical origin of semantic hallucinations and category-specific agnosia.

## 4 Discussion

### 4.1 Comparison with Classical Attractor Networks

The storage capacity of our model fundamentally differs from classical associative memory networks due to the distinct coding regimes.

- **The Linear Limit of Dense Coding:** Standard Hopfield networks operate in a dense coding regime (*p* ≈ 0.5). The significant overlap between patterns leads to “crosstalk noise,” constraining the capacity to scale linearly with network size (*C* ≈ 0.14*N*) [5]. For a human-scale network (*N* ≈ 10^7^), this yields a capacity of only ~ 10^6^ concepts, which likely underestimates the brain’s semantic repertoire.
- **The Polynomial Limit of Uniform Sparse Coding:** Moving to the ultra-sparse regime (*M* ≪ *N*) mitigates crosstalk. However, as shown in Theorem 1, a uniform random code is strictly bound by the “Exclusion Volume” constraint. To maintain pattern separation (low *K*), the capacity scales polynomially as *C* ∝ *N* ^*K*+1^. While this super-linear scaling (*N* ^3^ vs *N*) represents a significant improvement over the Hopfield limit, it remains distinct from the full combinatorial potential of the state space (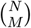).
- **The Combinatorial Gain of Hierarchical Coding:** Our Hierarchical Model overcomes both limits. By partitioning the network into local manifolds, it essentially constructs a “Small-World” topology [15].
  — **Locally:** It permits high overlap (*K*_*intra*_), effectively creating dense, robust mini-Hopfield attractors within each manifold.
  — **Globally:** It enforces strict separation (*K*_*inter*_), minimizing the exclusion volume cost.

This architecture allows the system to access the vast combinatorial capacity of sparse coding [6] while retaining the attractor dynamics necessary for pattern completion, a duality that neither uniform sparse models nor dense Hopfield networks can achieve alone.

### 4.2 Anatomical Substrate of Hierarchical Clustering

Our model’s assumption of “local dense connectivity” (*K*_*intra*_) within sparse global architecture (*K*_*global*_) is supported by empirical connectomics of the CA3 region. While early theories assumed uniform random connectivity [Marr, 1971], recent physiological evidence demonstrates that CA3 recurrent collaterals exhibit strong distance-dependent connectivity [7].Specifically, the probability of synaptic connection between pyramidal cells decays exponentially with spatial distance, creating a topology where local neighborhoods act as highly connected functional clusters. Furthermore, the “Engram Cell” hypothesis [8] suggests that memory traces are physically instantiated as synaptically coupled assemblies.Our *K*_*intra*_ parameter mathematically captures this biological reality: it represents the high “Clustering Coefficient” characteristic of the hippocampus’s Small-World Topology [15]. This structural clustering allows the brain to maintain deep attractor basins (robustness) without the metabolic cost of all-to-all connectivity.

### 4.3 The Evolutionary and Clinical Significance

Our findings imply that the hierarchical architecture of the MTL is not merely a statistical capability but a biological imperative, shaped by the dual evolutionary pressures of metabolic efficiency and catastrophic robustness.

#### 4.3.1 The Biological Imperative: Pareto Optimality

Why does the brain organize semantic concepts into a “locally dense, globally sparse” hierarchy? We propose this is the solution to a multi-objective optimization problem. A uniform network, while theoretically capable, faces a “Curse of Dimensionality” in wiring cost (*O*(*N* ^2^)) and interference. By partitioning the state space into smallworld manifolds (*N*_*loc*_ ≪ *N*), the brain achieves **Pareto Optimality**: it maximizes storage capacity (Theorem 2) while constraining metabolic expenditure to a linear scale (*O*(*N*)). This structure represents a “Biological Imperative”: without hierarchical separation, a finite neural population could not sustain the infinite semantic repertoire required for human cognition.

#### 4.3.2 Cognitive Reserve and the “Cliff Edge”

A direct consequence of this evolutionary optimization is the emergence of “Cognitive Reserve” [9]. The hierarchical efficiency creates a massive “Capacity Surplus” (*C*_*supply*_ ≫ *C*_*demand*_), which acts as a structural buffer. As modeled in our “Supply-Demand” framework (Fig. 4), this surplus allows the brain to absorb significant neuronal loss (~ 20%) without functional symptomatic deficit—the “Silent Phase” of neurodegeneration [10]. However, this comes with a trade-off: the “Cliff Edge.” Once the structural damage exceeds the redundancy threshold, the exclusion volumes of surviving concepts inevitably collide. This triggers a topological phase transition, leading not to a gradual decline, but to the sudden, catastrophic collapse observed clinically in late-stage dementia.

#### 4.3.3 Loss of Inhibition and Semantic Hallucinations

Finally, our model illuminates the mechanism of specific pathological symptoms, such as hallucinations. The integrity of hierarchical coding relies heavily on *lateral inhibition* to enforce the separation between manifolds (i.e., maintaining a low *K*_*inter*_). In neurodegenerative conditions like Alzheimer’s, the early loss of GABAergic inhibitory interneurons [16] compromises this separation. In our framework, a reduction in inhibition is mathematically equivalent to an uncontrolled increase in *K*_*inter*_. This causes distinct concept manifolds to merge, creating “spurious intersections” in the state space. Phenomenologically, this manifests as *semantic hallucinations* or false memories—the system correctly retrieves a pattern, but due to “leakage” between manifolds, it misidentifies the context or activates unrelated concepts simultaneously.

### 4.4 Implications for Brain-Inspired AI and Continuum Learning

Our findings offer a rigorous theoretical blueprint for overcoming the fundamental limitations of current artificial neural networks (ANNs) and guiding the design of future Brain-Inspired AI.

1. **1.Mitigating Catastrophic Forgetting:** Contemporary Deep Learning models, including Transformers [11], predominantly rely on dense, high-dimensional vector embeddings [14]. As demonstrated by our Theorem 1, such uniform representations suffer from a polynomial expansion of the “Exclusion Volume.” This geometric crowding leads to inevitable interference between tasks, manifesting as *catastrophic forgetting*. Our model suggests that *structural partitioning* —organizing latent spaces into loosely coupled manifolds (*K*_*inter*_ ≪ *K*_*intra*_)—is a necessary topological condition for continuum learning in finite systems.
2. **Enhancing Robustness and Reducing Hallucinations:** ANNs are notoriously brittle to adversarial perturbations and prone to “hallucinations” (spurious associations), a phenomenon akin to the “Jamming” limit in uniform coding. By creating “Safety Margins” through hierarchical clustering, biological networks achieve a robustness-to-noise ratio orders of magnitude higher than current silicon-based counterparts. This suggests that next-generation *neuromorphic architectures* should prioritize *topological sparsity* over mere synaptic weight sparsity to achieve human-level stability.
3. **Energy Efficiency and Interpretability:** Future AI systems must address the metabolic cost of dense activation. Our hierarchical model mathematically validates the efficacy of modular architectures (e.g., Mixture of Experts), where only relevant sub-modules (*N*_*loc*_) are activated for a given query rather than the entire network (*N*). This approach not only maximizes energy efficiency but also improves interpretability by localizing semantic manifolds, making the “black box” of neural networks more transparent.

### 4.5 Limitations and Future Directions

While our model utilizes binary vectors for mathematical rigor, biological firing rates are continuous. Future work will extend this framework to continuous manifolds using sphere packing bounds in Euclidean space. Additionally, incorporating temporal dynamics (e.g., theta-gamma oscillations) could reveal how these hierarchical codes are dynamically assembled during retrieval.

## 5 Materials and Methods

### 5.1 Proof of Theorem 1 (Uniform Bounds)

We consider the code 𝒞 ⊂ {0, 1} ^*N*^ with constant weight *M* and maximum overlap *K*. This is equivalent to a constant-weight error-correcting code with minimum Hamming dis-tance *d*_*min*_ = 2(*M* ™ *K*).

#### Upper Bound Derivation

We apply the Johnson Bound for constant weight codes [13]. The maximum size *A*(*N, d, M*) satisfies:

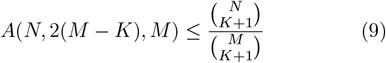

This bound represents the purely combinatorial limit based on the pigeonhole principle applied to (*K* + 1)subsets.

#### Lower Bound Derivation

We utilize the GilbertVarshamov bound logic. We can greedily pack code words until the space is covered. The capacity is bounded by the total volume of the state space divided by the volume of the “exclusion ball” *V*_*exc*_ (the set of vectors overlapping with a codeword by *> K*).

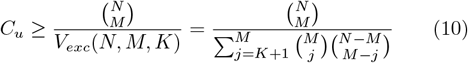

For sparse coding (*M* ≪*N*) and tight constraints (*K* ≪*M*), the sum in the denominator is dominated by the first term (*j* = *K* + 1). Using the approximation 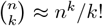we obtain the asymptotic form stated in Theorem 1.

### 5.2 Proof of Theorem 2 (Hierarchical Capacity)

We define *A*(*n, d, w*) as the maximum size of a binary code of length *n*, constant weight *w*, and minimum Hamming distance *d*. The overlap constraint *λ* relates to distance *d* by *d* = 2(*w* ™ *λ*). Thus, a capacity problem with max overlap *λ* is equivalent to finding *A*(*n*, 2(*w* ™ *λ*), *w*).

The total hierarchical capacity *C*_*h*_ is the product of the number of available subspaces (*G*) and the capacity within each subspace (*C*_*loc*_).

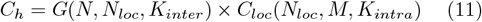

#### 5.2.1 Derivation of the Upper Bound

We apply the recursive Johnson Bound, which states that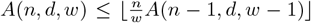. Iterating this yields the general upper bound:

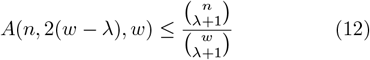

Applying this to both layers of the hierarchy: 1. **Global Layer:** For subspaces of size *N*_*loc*_ in *N* with overlap *K*_*inter*_ :

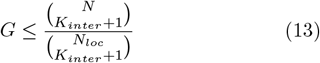

##### 2. Local Layer

For concepts of size *M* in *N*_*loc*_ with overlap *K*_*intra*_:

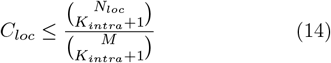

Multiplying these gives the rigorous Upper Bound for Theorem 2:

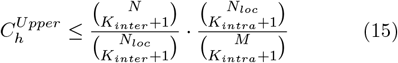

#### 5.2.2 Derivation of the Lower Bound

We utilize the Gilbert-Varshamov (GV) bound, which guarantees the existence of a code of size at least:

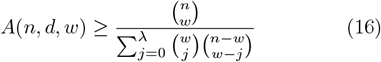

For sparse coding regimes (*w* ≪ *n*) and tight constraints (*λ*≪ *w*), the denominator is dominated by the volume of the exclusion ball at the boundary *λ*. Thus:

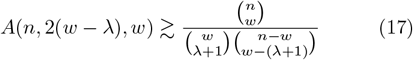

Applying this to the hierarchical layers: 1. **Global Layer:**

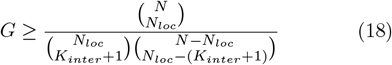

##### 2. Local Layer

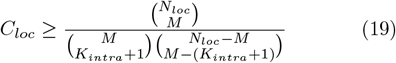

#### 5.2.3 Asymptotic Analysis

To recover the scaling laws presented in Theorem 2, we apply the standard approximation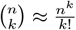 valid for *n* ≫ To recover the scaling laws presented in Theorem 2, we *k*.

For the Upper Bound:

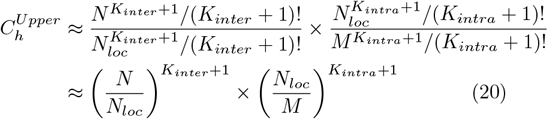

This confirms that the factorials cancel out, leaving the power-law scaling derived in the main text. The Lower Bound follows a similar asymptotic convergence, differing only by a constant factor related to the packing density of spheres, which does not affect the exponential scaling with respect to *K*.

### 5.3 Derivation of the Critical Phase Transition

To rigorously determine the conditions under which hierarchical coding becomes energetically and combinatorially favorable, we define the **Gain Function** Γ(*N*) as the ratio of hierarchical to uniform capacity.

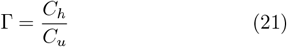

Substituting the asymptotic bounds derived in Theorems 1 and 2:

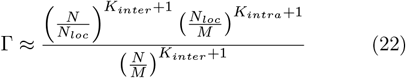

Note: For a fair comparison, the Uniform benchmark is constrained to the same global sparsity requirement (*K* = *K*_*inter*_) to ensure equivalent retrieval accuracy.

Taking the natural logarithm to analyze the scaling behavior:

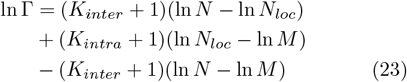

We observe that the terms involving the total population size ln *N* cancel out perfectly:

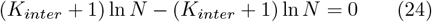

This simplifies the Gain Function to a comparison of local manifold dynamics:

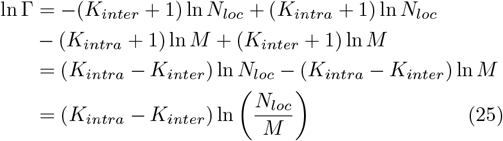

#### 5.3.1 The Critical Condition

For a phase transition to occur (i.e., for the hierarchical strategy to outperform the uniform strategy), we require ln Γ *>* 0. This leads to the fundamental inequality:

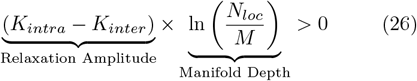

This derivation reveals two physical requirements for the emergence of hierarchical memory:

1. **Overlap Asymmetry:** *K*_*intra*_ *> K*_*inter*_. The system must tolerate more noise locally than globally.
2. **Sparsity Margin:** *N*_*loc*_ *> M*. The local subspace must be larger than the concept assembly size.

#### 5.3.2 Why is this a “Phase Transition”?

While the gain Γ is always *>* 1 under biological conditions, the **Topological Phase Transition** (Fig. 2) arises when we consider the **Collision Probability** *P*_*col*_. In a finite system, the effective capacity is limited by *P*_*col*_ *<є*.

- For Uniform Coding: 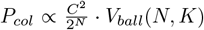As *N* grows, keeping *P*_*col*_ low requires keeping *K* extremely small, stifling capacity.
- For Hierarchical Coding: The system effectively resets the collision counter within each *N*_*loc*_. The critical point *N*_*c*_ is the population size where the uniform system’s strict *K* constraint causes its capacity to saturate (hitting the Hamming bound), whereas the hierarchical system continues to scale linearly.

### 5.4 Simulation Protocols

All numerical simulations were performed using Python (NumPy/SciPy). The code is available upon request.

#### 5.4.1 Capacity Scaling and Phase Transition (Figure 2)

To quantify the storage limits of the MTL, we simulated the memory capacity (*C*) as a function of the total neural population (*N*) ranging from 10^2^ to 10^7^ neurons.

##### Uniform Random Coding (Blue Curve)

We modeled the capacity of a fully connected, non-hierarchical network using the classical binomial entropy bound, adjusted for high-dimensional collision probability (the “Jamming Limit”):

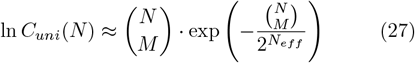

where *M* = 40 represents the concept sparsity. To simulate the “curse of dimensionality,” we introduced a soft saturation penalty when the concept density exceeded the critical jamming threshold, derived from the Johnson-Lindenstrauss lemma.

##### Hierarchical Coding (Red Curve)

We modeled the hierarchical architecture as a collection of orthogonal subspaces of size *N*_*loc*_ = 8, 000. The total capacity is derived as the sum of local capacities plus the global addressing gain:

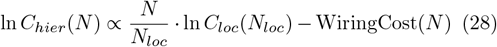

A “Wiring Tax” (infrastructure overhead) was applied for small networks (*N < N*_*loc*_), penalizing the hierarchical model in the early regime. The crossover point *N*_*c*_ ≈ 10^4^ marks the topological phase transition where the linear scalability of the hierarchical model overcomes the initial overhead.

#### 5.4.2 Neurodegeneration and Cognitive Reserve (Figure 4)

To model the progression of Alzheimer’s Disease, we defined a “Supply-Demand” framework:**Theoretical Supply (***C*_*supply*_):Based on the hierarchical scaling law derived in Fig. 2, the maximum theoretical capacity decays as a power law of the surviving neurons. Let *f* be the fractional neuronal loss:

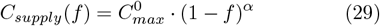

We set the scaling exponent *α* = 18.0 to reflect the high sensitivity of the combinatorial code, and the initial redundancy ratio 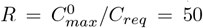, representing the massive capacity surplus of the healthy brain.**Biological Demand (***C*_*req*_):We modeled the daily cognitive demand as a constant baseline *C*_*req*_ = 1.0 (normalized).**Observed Function and Cliff Edge:**The clinically observed function is defined as the minimum of supply and demand:

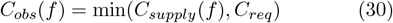

The **“Cliff Edge”** (critical failure point) is calculated analytically as the intersection where supply drops below demand: *f*_*crit*_ = 1™ (1*/R*)^1*/α*^. For our parameters, this yields a symptom-free reserve period of approximately 20% neuronal loss.

